# Elevated polygenic burden for ASD is associated with the broad autism phenotype

**DOI:** 10.1101/838375

**Authors:** K. Nayar, J.M. Sealock, N. Maltman, L. Bush, E.H. Cook, L.K. Davis, M. Losh

## Abstract

**Background:** Autism spectrum disorder (ASD) is a multifactorial, neurodevelopmental disorder that encompasses a complex and heterogeneous set of traits. Subclinical traits that mirror the core features of ASD, referred to as the broad autism phenotype (BAP) have been documented repeatedly in unaffected relatives and are believed to reflect underlying genetic liability to ASD. The BAP may help inform the etiology of ASD by allowing the stratification of families into more phenotypically and etiologically homogeneous subgroups. This study explored polygenic scores related to the BAP.

**Methods:** Phenotypic and genotypic information were obtained from 2,614 trios from Simons Simplex Sample. Polygenic scores of ASD (ASD-PGS) were generated across the sample to determine the shared genetic overlap between the BAP and ASD. Maternal and Paternal ASD-PGS was explored in relation to BAP traits and their child ASD symptomatology.

**Results:** Maternal pragmatic language was related to child’s social communicative atypicalities. In fathers, rigid personality was related to increased repetitive behaviors in children. Maternal (but not paternal) ASD-PGS was related to the pragmatic language and rigid BAP domains.

**Conclusions:** Domain- and sex-specific associations emerged between parent and child phenotypes. ASD-PGS associations emerged with BAP in mothers only, highlighting the potential for a female protective factor, and implicating the polygenic etiology of ASD-related phenotypes in the BAP.

## Introduction

Autism spectrum disorder is a neurodevelopmental disorder that is estimated to occur in 1 in 59 children below the age of 8 in the United States^1^. In their pioneering twin study, Folstein and Rutter^2^ evaluated monozygotic (i.e., identical) and dizygotic (i.e., fraternal) twins for infantile autism, finding higher concordance for infantile autism (i.e., both twins had the same diagnosis) in monozygotic twins (36%) than dizygotic twins (0%) and suggesting a genetic, heritable basis for ASD. A follow-up study^3^ using an expanded sample, revealed that 92% of monozygotic twin pairs were concordant for a *broader* spectrum of related social and abnormal cognitive atypicalities (compared to only 10% concordance in dizygotic pairs), suggesting that ASD and ASD traits were highly heritable. Indeed, a recent meta-analysis of ASD twin studies^4^ presented almost perfect concordance of ASD between monozygotic twins (98%), adding to the evidence of strong heritability of ASD. A substantial body of work has since identified a subclinical set of traits that mirror the core symptoms of ASD in unaffected relatives, including social reticence, rigid personality, and pragmatic (i.e., social) language differences, collectively known as the broad autism phenotype (BAP).

Component features of the BAP may constitute candidate endophenotypes, or heritable traits linked to a disorder that are observed in affected individuals and unaffected relatives. Though many endophenotypes are likely polygenic themselves, they nonetheless may be useful for study in complex, heterogeneous conditions such as ASD, by helping to identify etiologically homogeneous subgroups based on shared endophenotypes^5–7^. Evidence that features of the BAP constitute candidate endophenotypes comes from studies showing significantly higher rates of BAP features in unaffected relatives of individuals with ASD than the general population^8–11^, and even higher in multiplex families^12,13^, suggesting that BAP features are sensitive indices of genetic loading. ASD symptomatology in children has also been shown to correlate with BAP features in parents^14–16^, even when BAP traits were noted before a parent went on to have a child with ASD^17^. Further, BAP features have been shown to cosegregate with distinct patterns of neurocognitive performance in clinically unaffected relatives^12,18–22^, suggesting links with underlying neural substrates impacted by ASD genetic risk. Finally, unlike ASD, which by definition requires the presence of impairments in social interaction and communication, and the presence of restricted and repetitive interests and behaviors, traits of the BAP have been observed to segregate independently in unaffected relatives^22–24^, potentially reflecting distinct genetic contributions to the component features of ASD^25–27^. Taken together, evidence suggests that the BAP may therefore help to leverage studies of ASD etiology by providing a distilled phenotypic expression of genetic liability to ASD with potentially more straightforward ties to underlying etiology than the complex and heterogeneous ASD phenotype^28^. Characterizing endophenotypes among clinically unaffected relatives may also provide insights into familial patterns of transmission, permitting focus on transmitting relatives for more refined analysis of ASD risk loci^11,28^.

### ASD polygenic etiology and scores

As noted previously, family studies of ASD show an aggregation of cases within families, suggesting an inherited genetic component^29,30^. Family and population genetic studies have revealed a complex architecture with genetic liability originating from rare, structural, *de novo*, and common variation^29,31^. However, rare, structural, and *de novo* variants collectively explain less than 5% of the total liability of autism^32,33^. In contrast, common variants are estimated to explain the majority of the genetic contribution to ASD^32,34^. Genome-wide association studies estimate that narrow sense heritability due to common variation is approximately 12% for ASD^35^. One method to assess associations with ASD genetics is by harnessing common variants through polygenic scoring.

Polygenic scoring is a method to calculate an individual’s underlying genetic liability to a complex trait using weights derived from large-scale genome-wide association studies. A major benefit of polygenic scores (PGS) is that a score can be calculated for any genotyped individual, regardless of diagnostic status. PGS of ASD have been previously shown to imperfectly, but significantly, predict ASD case-control status and ASD-related features^34^, suggesting reliable measurement of inherited ASD genetic factors. In addition to a high degree of sensitivity in diagnostic prediction, the range of derived PGS allows for continuous measurement of use in correlational analyses with other complex phenotypes. For example, ASD-PGS associate with general cognitive ability, logical memory, and verbal intelligence in the general population^36^. Finally, given the polygenic nature of ASD with high levels of locus of heterogeneity, together with heterogeneous symptom presentation, studying inherited variants involving a large number of genes, such as ASD-PGS, in family members of individuals with ASD can help to increase power to identify the transmission of polygenic risk into the next generation, as opposed to relying on the low frequency single candidate gene linkages (although, another fruitful approach is usage of genome-wide linkage analysis)^28^.

This study examined PGS within families of individuals with ASD, with the hypothesis that ASD-PGS underlie both aggregate ASD expression and subclinical BAP features. It was predicted that those child measures that are significantly associated with parent BAP scores would also emerge as an important phenotype in subsequent ASD-PGS analyses. Specifically, the study aimed to examine 1) how ASD-PGS relate to BAP features in parents of individuals with ASD; 2) how ASD-PGS relate to phenotypic variation in individuals with ASD; and 3) the familiality of ASD-PGS and associated phenotypes, by assessing parent-child phenotypic and ASD-PGS associations.

## Methods and Materials

### Participants

Participants were drawn from the Simons Simplex Collection (SSC) and included a maximum of 2,618 mothers, 2,614 fathers, and 2,621 children with ASD. Inclusion criteria for SSC required that probands were aged between 4-18 years, had non-verbal mental age above 18 months, no history of neurological deficits, birth trauma, perinatal complications, genetic evidence of fragile X syndrome or Down syndrome^37^, or ASD within 3^rd^ degree relatives, as well as inclusion of both parents when available. All children included in the study met DSM-IV or DSM-5 diagnostic criteria for ASD. The child sample included 348 females and 2280 males (n = 13 sex missing; note that these individuals were included in phenotypic analyses but were removed from genetic analyses). Parent and child participant characteristics are further described in *Table 1*.

**Table 1:**
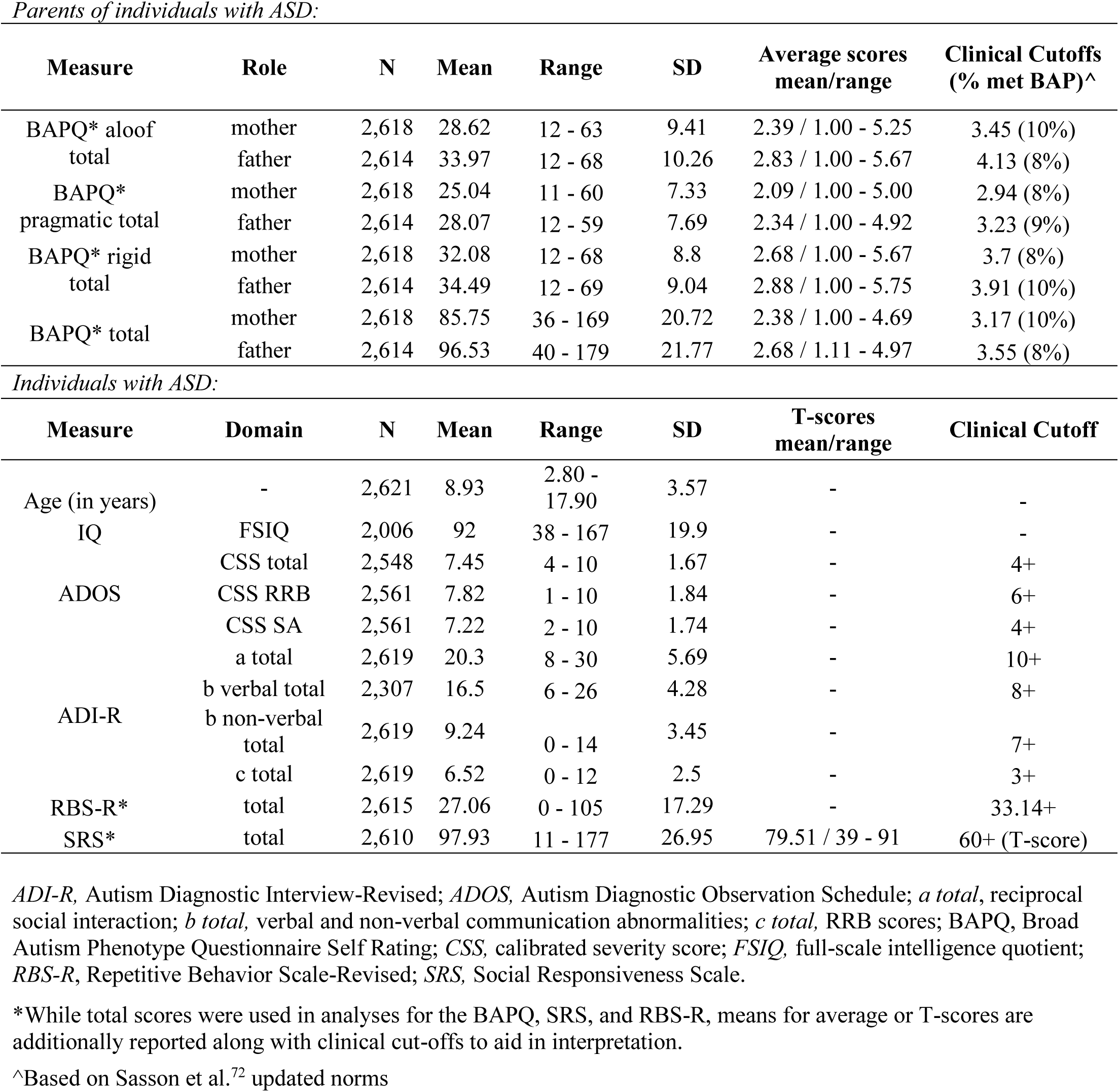
Sample characteristics

### Phenotypic Measures

#### Data quality control

Parent and child phenotypes were derived based on clinical assessment and questionnaire measures aimed at characterizing BAP and ASD traits, respectively. Given the large sample size of the SSC, as well as the number of different measures included in the repository, a series of *a priori* analyses were completed to determine those measures that best captured a range of ASD-related traits within the constraints of a continuous distribution for PGS predictive analyses. Distributions for each measure were examined separately for normality and to detect potential outliers. For each parent measure, distributions were explored with all parents combined, as well as for mothers and fathers separately. For each child measure, distributions were similarly examined, though sex remained collapsed due to the smaller female sample. Parent-child correlations were examined to determine whether child phenotypes related to BAP traits in parents.

### Parent BAP Measures

The Broad Autism Phenotype Questionnaire (BAPQ)^39^ is a 36-item questionnaire that assesses the personality and pragmatic language traits associated with the core deficits of ASD, and which are known to be associated with the Broad Autism Phenotype (BAP). As described above, distributions for each measure were examined for normality to ensure that assumptions for parametric analyses were met. Patterns of sex differences for parents of children with ASD emerged across BAPQ subscales and total scores, consistent with prior findings^38,39^. Mothers and fathers were therefore examined separately for all analyses. Both BAPQ total score, and the subdomain scores for Aloof, Pragmatic Language, and Rigid were examined.

### Proband ASD Measures

The following measures were explored in relationship with paternal and maternal BAPQ subscale and total scores: i) Autism Diagnostic Observation Schedule (ADOS) calibrated severity scores (CSS)^40,41^; ii) Autism Diagnostic Interview-Revised (ADI-R)^42^; iii) Repetitive Behavior Scale-Revised (RBS-R)^43^; iv) Social Responsiveness Scale (SRS)^44^ parent report^45^; v) Social Communication Questionnaire (SCQ)^46^; vi) Aberrant Behavior Checklist (ABC)^47^; vii) Child Behavior Checklist (CBCL)^48^; and viii) Vineland Adaptive Behavior Scales (VABS)^49^. The SCQ, ABC, CBCL and VABS were dropped from subsequent analyses given the lack of associations observed between these measures and parent BAPQ, and in order to reduce the number of variables contributing to the regression models (specific subscores included are outlined in Table 1).

As part of the SSC data collection procedures, different IQ assessments were administered to participants based on age and cognitive ability. Assessments with the largest sample size were prioritized, followed by those with lower sample sizes: Wechsler Intelligence Scale for Children (WISC), followed by the Wechsler Abbreviated Scale of Intelligence (WASI), followed by the Differential Ability Scales *School-Age* (DAS-SA), followed by Differential Ability Scales *Early Years* (DAS-EY). To compute full-scale IQ, the PRI and VCI were included for the WISC, VIQ and PIQ were included for the WASI, and Special Nonverbal Composite (SNC) and Verbal scores were included for the DAS-SA and DAS-EY (note: given that the DAS EY *Upper Level* included an SNC score while the *Lower Level* included a non-verbal reasoning score, a combined SNC score was generated to equate the two). There was no minimum or maximum IQ set for these analyses, particularly given the heterogeneity observed in cognitive abilities in ASD^50^.

### Genetic Data Quality Control

SSC samples were genotyped on one of three Illumina platforms, 1Mv1, 1Mv3 or Omni2.5. Quality control was performed on each of the platforms separately using PLINK v1.9^51^. First, SNPs were filtered at a call rate of 0.95. Individuals were filtered if their genotype missingness rate was greater than 0.02, heterozygosity above 0.2 or below −0.2, or if there were discrepancies between the number of sex chromosomes and reported sex. Next, SNPs were filtered more stringently for call rates below 0.98 and differential missingness above 0.02. We then used LiftOver^52^ to convert the platforms from genome build hg18 to hg19 to match the 1000 Genomes Project^53^ sample and the summary statistics used for polygenic scoring. Genotype Harmonizer^54^ was used to align SNPs with strand information from 1000 Genomes Project Phase 3. We performed principal component analysis on parental controls using Eigenstrat^55,56^. To check for platform effects, we performed logistic regressions between each platform using controls and covarying for top 10 principal components of ancestry and sex. We removed SNPs significantly associated with platform (*p* < 0.001). To define the European individuals, we performed PCA with a combined sample of self-reported white parents and the 1000 Genomes Project Phase 3 sample, and selected individuals around the CEU cluster. For probands and siblings to be included, both parents were required to be of European genetic ancestry. Finally, we filtered the European parental sample for SNPs out of Hardy-Weinberg Equilibrium (*p* < 1e^-6^) and removed the same SNPs from probands and siblings.

### Polygenic Score Generation

For polygenic scoring, we obtained summary statistics from the Psychiatric Genetics Consortium’s meta-analysis of Autism^35^ after removing the SSC sample. Polygenic scores (PGS) were generated using PRScs-auto^57^, using the CEU sample from 1000 Genomes as the LD reference panel. PRScs uses a Bayesian framework to model linkage disequilibrium from an external reference and applies a continuous shrinkage prior on SNP effect sizes to adjust for linkage disequilibrium. Using PLINK v1.9 with the PRScs adjusted summary statistics, we generated polygenic scores on each platform separately. Polygenic scores were calculated on 1,018 individuals on platform 1Mv1 using 789,419 SNPs, 3,253 individuals on 1Mv3 with 789,419 SNPs, and 2,983 individuals on Omni2.5 with 627,649 SNPs. To analyze all three platforms together, we adjusted the PGS for platform effects and z-score scaled the residuals (Supplementary Figure 1).

### Statistical Analysis

A series of linear regression analyses were conducted using base-R or the lmer package^58^ for R. Details of regression statistical analyses are outlined below.

#### Parent-child phenotype associations

All measures reported above were included in phenotypic analyses with sample characteristics and sizes detailed in *Table 1.* For all phenotype-phenotype analyses, *p*-values were stringently corrected using the Bonferroni method to account for 72 tests performed (*p* = .0007). Each parent-child association was explored for mothers and fathers separately. Despite significant associations between father and mother BAPQ scores and child IQ (*p*s < .01), no covariates were added to the regression model^50^.

#### Genotype-phenotype analyses

All phenotypic data reported above were used for polygenic analyses, however sample sizes differed by measure as follows: BAPQ aloof, pragmatic, rigid, and total scores (n_mothers_ = 1,812, n_fathers_ = 1,808), ADOS total CSS (n = 1,765), ADOS RRB CSS (n = 1,782), ADOS SA CSS (n = 1,782), ADI a total (n = 1,826), ADI b nonverbal total (n = 1,826), ADI b verbal total (n = 1,644), ADI c total (n = 1,826), RBS-R total (n = 1,824), SRS total (n = 1,827), and IQ scores (n = 1,443). All scores were z-score scaled for analysis so that the odds ratios were per 1 SD increase in PGS.

#### Parental BAPQ Phenotypes and Parental Polygenic Scores

The association between parental PGS and parental BAPQ phenotypes was explored using a set of linear regressions between maternal or paternal PGS and maternal or paternal BAPQ phenotypic scores, respectively, adjusting for top 10 principal components of ancestry. Statistical significance was determined separately for mothers and fathers using Bonferroni adjustment (*p* < 0.0125).

#### Proband Phenotypes and Proband Polygenic Scores

Linear regressions between proband PGS and proband phenotypes were performed, adjusting for proband sex and top 10 principal components of ancestry. Statistical significance was set at *p* < 0.005.

#### Parental BAPQ Phenotypes and Proband Polygenic Scores

Associations between parent phenotypes and proband PGS were examined by parent sex, which involved linear regressions between maternal/paternal BAPQ scores and the proband PGS. Statistical significance was determined separately for mothers and fathers using Bonferroni adjustment (*p* < 0.0125).

#### Associations Between Proband Phenotypes and Parental Polygenic Scores

To determine associations between proband phenotypes and parent PGS, a set of linear regressions were examined between each proband phenotype and maternal and paternal PGS, separately. Statistical significance was determined separately for mothers and fathers using Bonferroni adjustment (*p* < 0.005).

## Results

### Parent-child phenotypic associations (*Fig. 1*)

#### ADOS

There were no significant associations between maternal or paternal BAPQ scores and proband ADOS total CSS or any of the ADOS subscales (SA CSS, RRB CSS).

#### ADI-R

A significant positive association was detected between maternal scores on the BAPQ-pragmatic subscale and proband ADI-R a (reciprocal social interaction) and b (non-verbal communication) total scores (*Estimates* > .09, *p*s < .0007) **(*Fig. 2a*).** There were no significant associations between paternal BAPQ scores or maternal BAPQ-aloof, -rigid, or -total scores and proband ADI-R scores.

**Figure 1.**
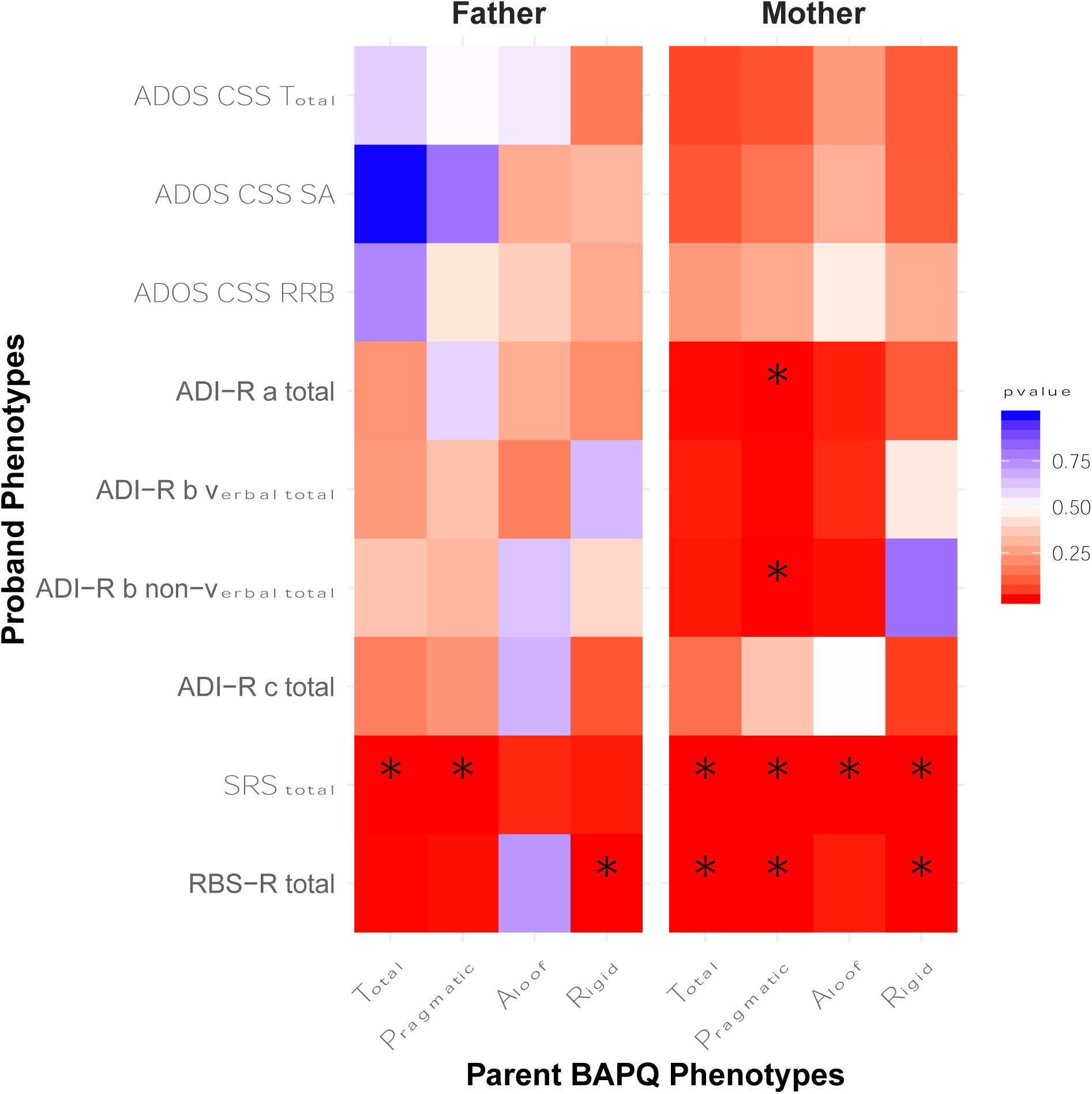
Heatmap of associations between paternal (left) and maternal (right) BAPQ scores and proband clinical-behavioral scores. Asterisks denote associations passing Bonferroni correction.

**Figure 2.**
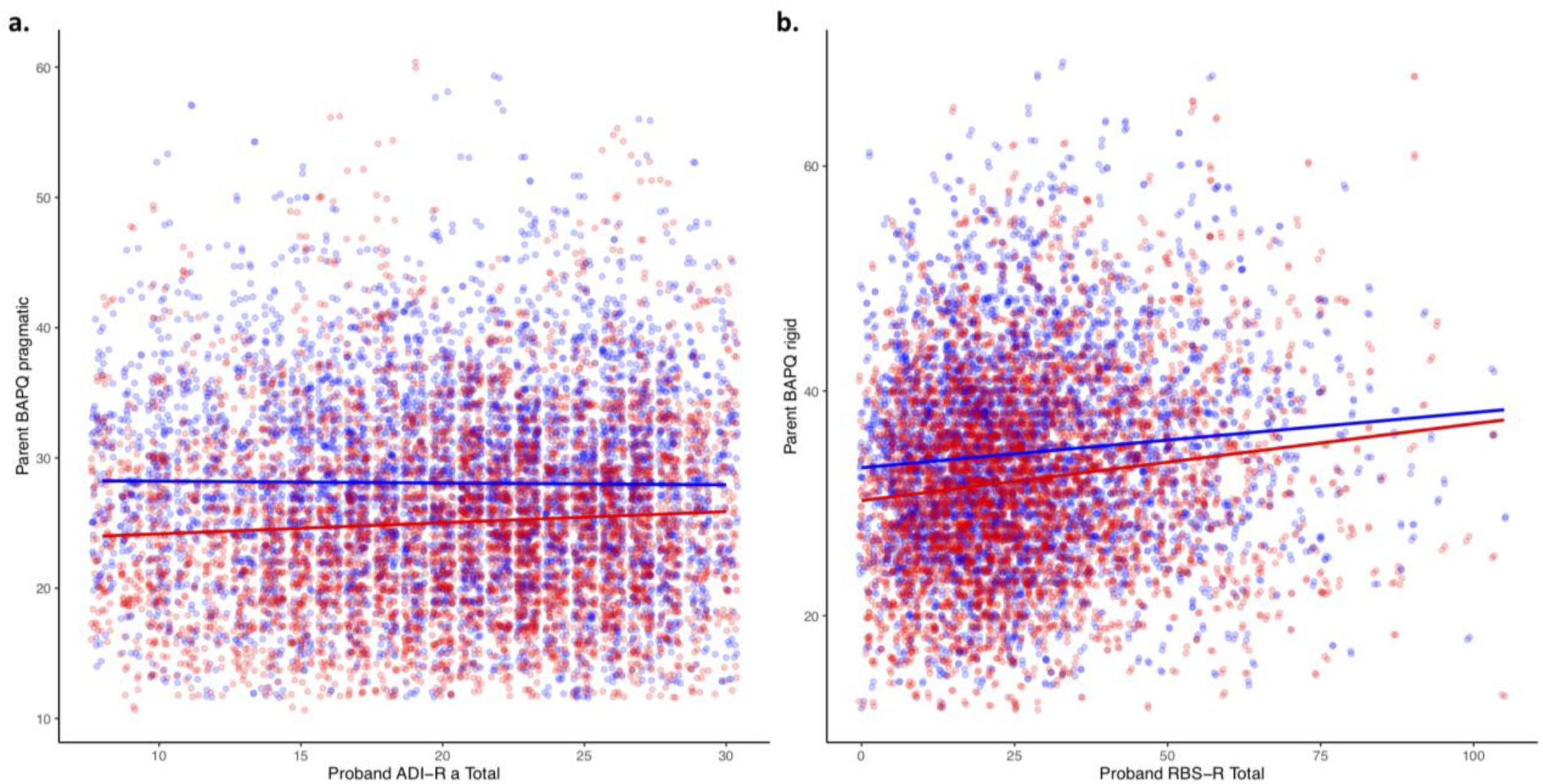
Relationship between proband ADI-R a total scores and parent BAPQ pragmatic scores (a); and proband RBS-R total scores and parent BAPQ rigid scores (b). Maternal regression line in red; paternal regression line in blue.

#### SRS

All mother BAPQ scores were significantly positively associated with proband SRS total scores (*Estimates* > .03, *p*s < .0007). In contrast, only paternal BAPQ-pragmatic and -total scores related to SRS-total scores in probands (*Estimate*s > .02, *p*s < .0007). No associations were observed between paternal BAPQ rigid or aloof and proband SRS scores.

#### RBS-R

Paternal BAPQ-rigid scores were significantly positively associated with proband RBS-R total scores (*Estimate* = .05, *p* < .0007) **(*Fig. 2b*)**. In contrast, mother BAPQ-pragmatic, - rigid, and -total scores were all associated with proband RBS-R total scores (*Estimates* > .06, *p*s < .0007). No significant association between paternal BAPQ-aloof, -pragmatic, or -total, or maternal BAPQ-aloof scores and proband RBS-R total scores emerged.

### Genotype-Phenotype associations (*Fig. 3, Supplementary Table 1*)

#### Parental BAPQ Phenotypes and Polygenic Scores (Fig. 4)

Maternal polygenic score were associated with BAPQ-pragmatic (OR = 1.09, *p* = 2.62 × 10^−4^, SE = 0.02), -rigid (OR = 1.06, *p* = 0.012, SE = 0.02), and -total (OR = 1.09, *p* = 5.13 × 10^−4^, SE = 0.02) scores. However, paternal ASD-PGS were not associated with any phenotypes tested. Additionally, the effect estimates of maternal PGS on maternal pragmatic-, rigid-, and total-BAPQ scores were increased compared to the effects of paternal PGS on paternal BAPQ scores (Supplementary Table 1).

**Figure 3.**
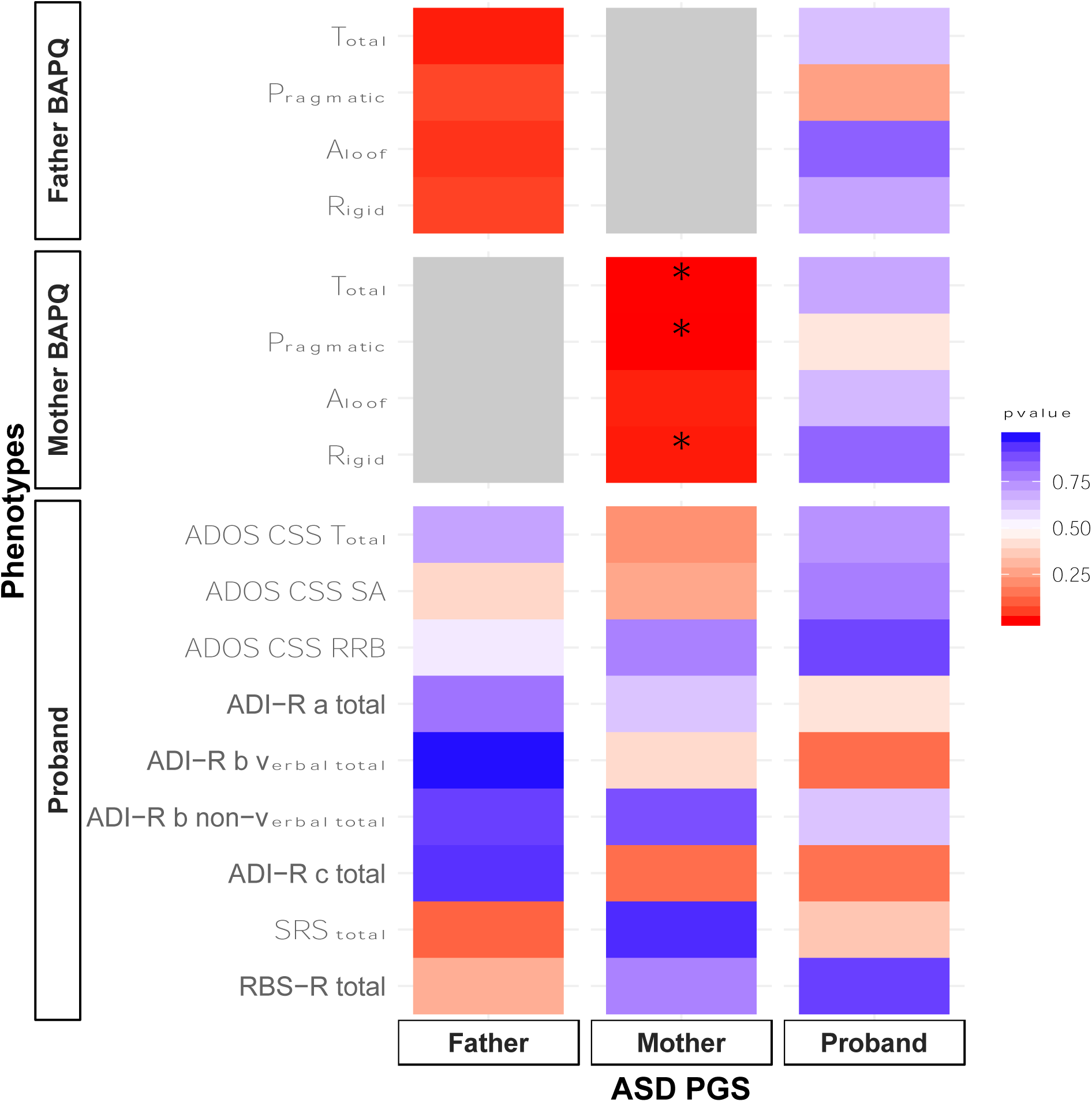
Heatmap of associations between paternal, maternal, and proband PGS with clinical-behavioral features of ASD and the BAP. Asterisks denote associations passing Bonferroni correction.

**Figure 4.**
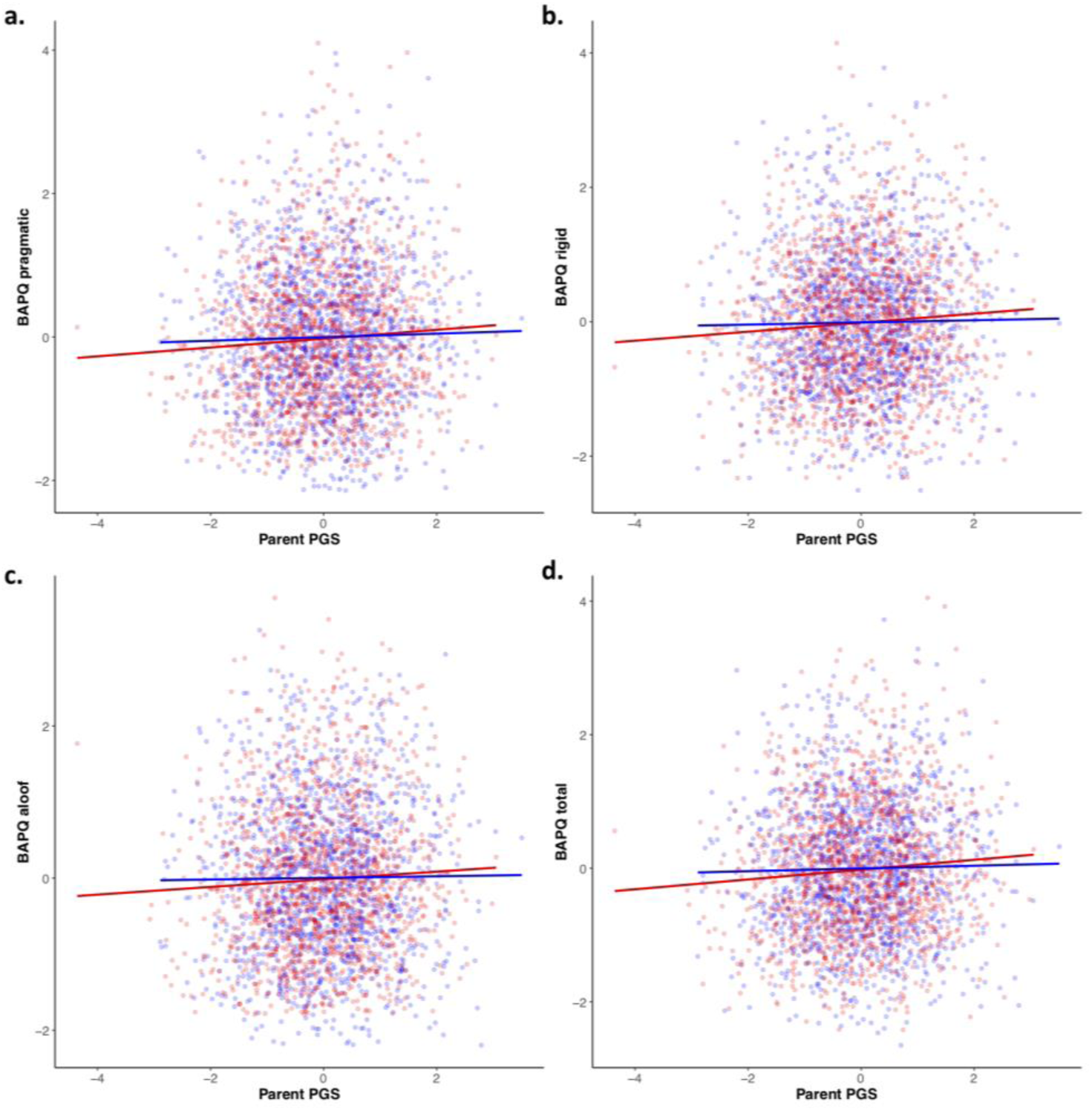
Relationships between maternal and paternal PGS of ASD, and BAPQ pragmatic (a), rigid (b), aloof (c), and total (d) scores. Mothers are plotted in red. Fathers are plotted in blue. Linear regression lines are plotted for mothers and fathers separately.

#### Proband Phenotypes and Proband Polygenic Scores

Proband ASD-PGS were not associated with any phenotypes tested.

#### Parental BAPQ Phenotypes and Proband Polygenic Scores

Proband ASD-PGS did not show any associations with mother’s or father’s BAPQ scores.

#### Proband Phenotypes and Parental Polygenic Scores

None of the tested proband phenotypes showed associations with maternal or paternal ASD-PGS.

## Discussion

This study examined polygenic scores (PGS) within families of individuals with ASD, with the goal of exploring parent and child genotype-phenotype and phenotype-phenotype associations, as well as familial polygenic liability associated with ASD. Overall, results demonstrated relationships between parent and child clinical-behavioral phenotypes, as well as associations between parents’ PGS and features of the broad autism phenotype (BAP). Additionally, a formal transmission disequilibrium test replicated a previous report^59^ and showed unequal polygenic transmission between probands and unaffected siblings (data not shown). Given the likelihood of increased *de novo* mutations in simplex families studied here, our results further emphasize the role of inherited genetic risk associated with ASD and the constituent features of ASD and the BAP.

Consistent with prior work, no definitive sex-differences emerged in BAP trait expression^60,61^. Nonetheless, numerous parent-child clinical-behavioral relationships emerged in this study, with robust parent of origin associations. In mothers, language-related phenotypes were consistently associated with more severe ASD symptomatology in probands. Specifically, maternal pragmatic language scores were strongly associated with children’s social and non-verbal communication skills measured by the ADI-R. Similar parent-child associations have been observed in other studies examining language-related phenotypes, where subtle differences in language fluency in mothers with the BAP were found to relate to more severe symptoms in their children with ASD^19^. Such a pattern of lineality may suggest a stronger inherited maternal effect for language-related phenotypes in ASD (though important to consider is that parents and children influence one another’s language patterns as well). Whereas the maternal effect appeared to be centered around language-related phenotypes, paternal BAP features appeared more associated with the RRB/rigid domain. In contrast, all domains of the BAP in mothers were related to RRBs in probands. Despite there being no prior family history of ASD in the families included in this study, these patterns of domain-specific familiality may be further evidence that constituent features related to ASD combine additively to increase ASD risk^62^.

In line with findings from phenotypic analyses within families, analyses of parents’ ASD-PGS also demonstrated a robust genetic effect on mothers’ phenotypes, such that polygenic variants associated with ASD predicted maternal BAP features, including pragmatic language differences and rigid personality. The lack of paternal associations is not likely to be due to a lack of power, since the same sample size in mothers yielded significant associations. Importantly, the effect estimates of maternal PGS on BAPQ scores was in some cases almost double the effect of paternal PGS on paternal BAPQ scores, which could indicate a sex-specific influence of ASD-risk genes on the BAP. Sex differences are well-documented in both diagnostic rates and phenotypic expression in males and females with ASD^63,64^, and have been hypothesized to result from a female protective effect^65^, where females require greater inherited risk than males to exhibit ASD. Although large-scale studies have shown no appreciable difference in the common variant liability between males and females with ASD^66^, studies have shown an enrichment of loss-of-function *de novo* variants and rare copy number variants in female cases compared to controls^67^. The sex-specific phenotypic and genotypic-phenotypic associations that emerged here may provide evidence for a female genetic protection for common variation as it relates to the ASD phenotypic spectrum. It is therefore possible that females require both increased ASD-PGS and rare variation to lead to ASD, while only ASD-PGS in females leads to the BAP; in contrast, the same PGS in males would lead to the clinical manifestation of ASD. The discrepancy between studies may be due to the generally smaller female ASD sample sizes included in prior work^63^ in comparison to the larger sample sizes of unaffected mothers with increased genetic liability to ASD included in the present study, likely increasing power to detect a female protective effect on domain phenotypes. As such, this study may inform understanding of familial transmission of ASD-related traits, which may help to enhance our power to detect genetic phenotypic variants associated with ASD.

Finally, the lack of associations between proband ASD-PGS or parental BAP scores with proband clinical-behavioral features, as well as between parental PGS and proband clinical-behavioral phenotypes, were somewhat surprising. Lack of associations may, however, be a reflection of the phenotypic homogeneity present in the SSC sample (given stringent selection criteria of the SSC), and/or potentially higher intellectual ability of parents in the SSC (though, not directly assessed in parents in the SSC sample), which has been found to be associated with greater polygenic risk^36,68–70^. Inclusion of only simplex families from SFARI Base SSC may additionally explain the lack of findings, with theorized additive genetic risk more commonly occurring in multiplex versus simplex families^71^. Furthermore, the SSC excluded families where ASD was suspected in parents, potentially reducing the variance of BAP features in the sample and reducing the ability to find associations with BAP features. As such, future studies should consider inclusion of multiplex families. Future work may also benefit from the inclusion of additional BAP assessments in analyses, as questionnaires may limit the sensitivity of BAP trait detection given potential reporting biases in self-report measures of BAP traits^38^.

In sum, this study revealed key trends toward sex-specific associations of ASD-related features in probands with ASD and the parental BAP, with effects on mothers emerging in the language-related social-communication domain, and paternal phenotypic effects in the rigid/RRB domain. Additionally, ASD-PGS associations with the BAP emerged only in mothers, highlighting the potential for a female protective factor that may also be expressed among first-degree relatives of individuals with ASD. Together, findings from this study underscore the significance of the BAP in parents, which may reflect more influences from common genetic variation and polygenic risk rather than rare variation that contributes to aggregate and heterogeneous ASD^26^.

## Acknowledgments

We are grateful to all of the families at the participating Simons Simplex Collection (SSC) sites, as well as the principal investigators (A. Beaudet, R. Bernier, J. Constantino, E. Cook, E. Fombonne, D. Geschwind, R. Goin-Kochel, E. Hanson, D. Grice, A. Klin, D. Ledbetter, C. Lord, C. Martin, D. Martin, R. Maxim, J. Miles, O. Ousley, K. Pelphrey, B. Peterson, J. Piggot, C. Saulnier, M. State, W. Stone, J. Sutcliffe, C. Walsh, Z. Warren, E. Wijsman). We appreciate obtaining access to phenotypic and genetic data on SFARI Base. Approved researchers can obtain the SSC population dataset described in this study (https://www.sfari.org/resource/simons-simplex-collection/) by applying at https://base.sfari.org. We also thank Dr. Richard Anney for providing the PGC ASD summary statistics excluding the SSC samples. The content is solely the responsibility of the authors and does not necessarily represent the official views of the National Institutes of Health. Research time was supported by grants from the National Institutes of Health (R01DC010191, PI: Losh). JMS is funded by NIH/NIGMS 5T32GM080178-12 (PI: Dr. Nancy J. Cox).

## Author contributions

ML, LKD, and ECH conceived and designed the study and oversaw all analyses, and writing of the manuscript. KN and JMS led data processing, analysis, and manuscript preparation. NM and LB contributed to data processing and analysis. NM also contributed to manuscript preparation. All authors read and approved the final manuscript.

## Conflict of Interests

Authors declare that they have no disclosures or conflicts of interest

---

**Supplementary Figure 1.**
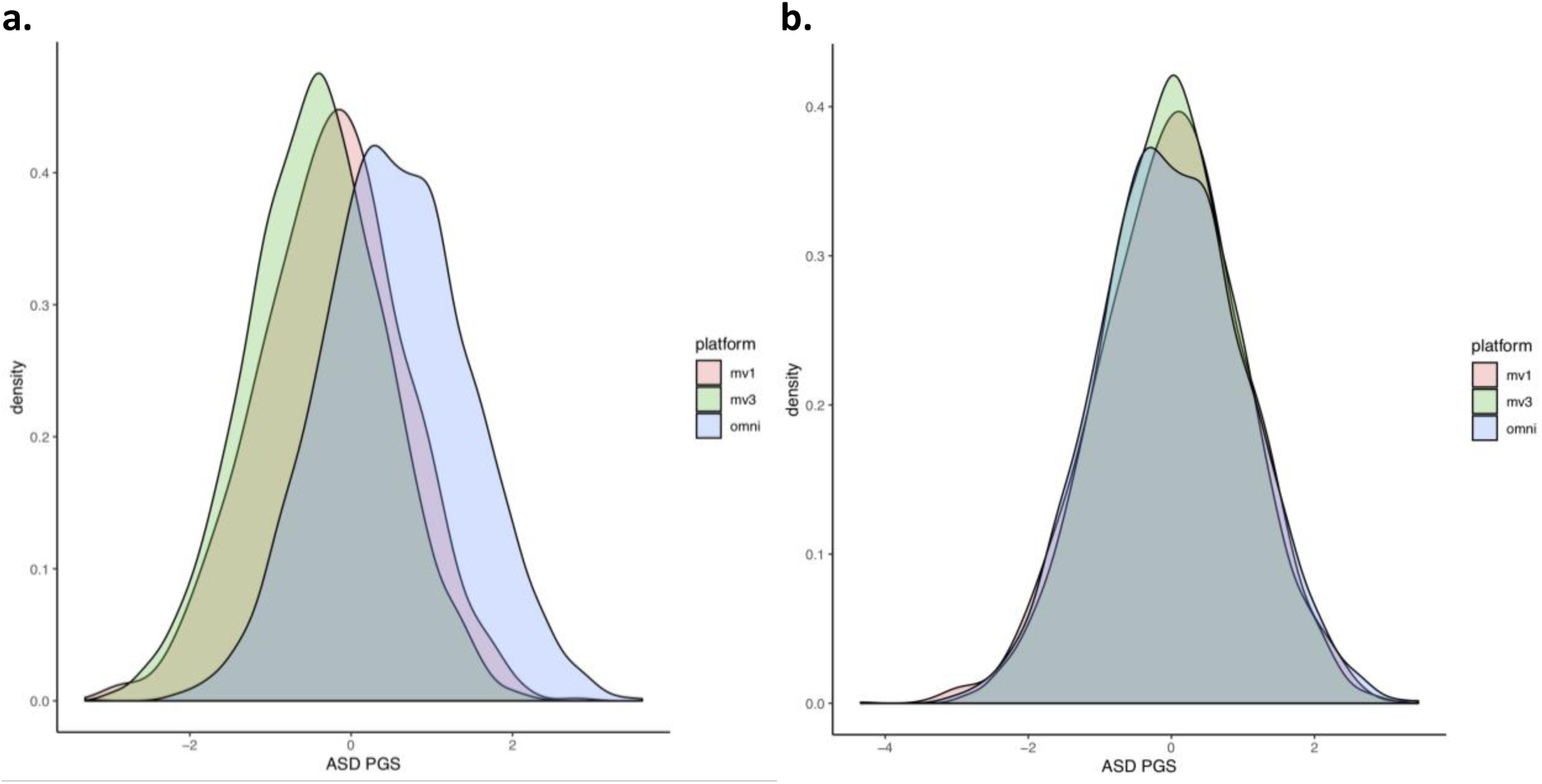
Distribution of ASD PGS color-coded by platform a) z-score scaled and b) residual of platform and z-score scaled.

**Supplementary Table 1.** Associations between parental ASD PGS and BAPQ phenotypes.

| **BAPQ Measurement** | **PGS** | **p-value** | **beta** | **SE** |
| --- | --- | --- | --- | --- |
| **Aloof** | Mother | 1.70E-02 | 0.06 | 0.02 |
|  | Father | 3.58E-02 | 0.05 | 0.02 |
| **Pragmatic** | Mother | 2.62E-04* | 0.09 | 0.02 |
|  | Father | 6.24E-02 | 0.05 | 0.02 |
| **Rigid** | Mother | 1.20E-02* | 0.06 | 0.02 |
|  | Father | 5.72E-02 | 0.05 | 0.02 |
| **Total** | Mother | 5.13E-04* | 0.09 | 0.02 |
|  | Father | 1.40E-02 | 0.06 | 0.02 |

\*Associations passing multiple testing correction

